# Stabilin-1 plays a protective role against *Listeria* monocytogenes infection through the regulation of cytokine and chemokine production and immune cell recruitment

**DOI:** 10.1101/2021.02.23.432451

**Authors:** Rita Pombinho, Jorge Pinheiro, Mariana Resende, Diana Meireles, Sirpa Jalkanen, Sandra Sousa, Didier Cabanes

**Affiliations:** Instituto de Investigação e Inovação em Saúde - i3S, Universidade do Porto, Porto, Portugal; Group of Molecular Microbiology, Instituto de Biologia Molecular e Celular - IBMC, Porto, Portugal; Microbiology and Immunology of Infection, Instituto de Biologia Molecular e Celular - IBMC, Porto, Portugal; MediCity Research Laboratory and Department of Medical Microbiology and Immunology, University of Turku, Turku, Finland

**Keywords:** *Listeria*, Scavenger Receptors, STAB-1, innate immunity, infection

## Abstract

Scavenger receptors are part of a complex surveillance system expressed by host cells to efficiently orchestrate innate immune response against bacterial infections. Stabilin-1 (STAB-1) is a scavenger receptor involved in cell trafficking, inflammation and cancer, however its role in infection remains to be elucidated. *Listeria monocytogenes* (*Lm*) is a major intracellular human food-borne pathogen causing severe infections in susceptible hosts. Using a mouse model of infection, we demonstrate here that STAB-1 controls *Lm*-induced cytokine and chemokine production and immune cell accumulation in *Lm*-infected organs. We show that STAB-1 also regulates the recruitment of myeloid cells in response to *Lm* infection and contributes to clear circulating bacteria. In addition, whereas STAB-1 appears to promote bacterial uptake by macrophages, infection by pathogenic *Listeria* induces the down regulation of STAB-1 expression and its delocalization from the host cell membrane.

We propose STAB-1 as a new SR involved in the control of *Lm* infection through the regulation of host defense mechanisms, a process that would be targeted by bacterial virulence factors to promote infection.

## Introduction

*Listeria monocytogenes* (*Lm*) is a major human food-borne pathogen that causes listeriosis, which is highly prevalent among high-risk groups including immunocompromised people, elderly, pregnant women and neonates. Listeriosis is an overall public health concern associated with high hospitalization and mortality rates, being the most deadly food-borne infection in Europe [1]. Manifestations of the disease range from a self-limiting febrile gastroenteritis to septicaemia, meningitis and encephalitis [2]. The most severe aspects of the disease are related to the capacity of *Lm* to cross the intestinal, blood-brain and maternal-foetal barriers, evading the immune response, multiplying within phagocytic and non-phagocytic cells and effectively disseminating throughout host tissues [3]. These properties are shaped by an arsenal of virulence factors [4].

The host innate immune response is critical to elicit an early defense towards *Lm.* The containment of infection requires both the participation of professional phagocytes that trap bacteria from target organs, and the activation of a number of pattern recognition receptors, including scavenger receptors (SRs) [5, 6]. SRs comprise a diverse and conserved family of proteins, able to bind to a wide range of ligands stimulating the removal of non-self and modified-self targets [7]. They contribute to maintain homeostasis and control pathogen infections, playing key functions in the antimicrobial host immune response [7, 8]. The role of SRs in *Lm* infection was first revealed for SR-A, SR-AI/II KO mice showing increased susceptibility to *Lm* infection and displaying increased hepatic granuloma formation [9]. Later, MARCO, CD36 and SR-BI were then shown to bind *Lm* and to modulate the immune response against *Lm* [10–12]. The first member of the Class H of SRs to be described was STABILIN-1 (STAB-1) [13]. It is a highly conserved type I transmembrane protein mainly expressed in sinusoidal endothelial cells of the spleen and liver, and on both afferent and efferent arms of the lymphatic vasculature, but also in subpopulations of monocytes/macrophages, and hematopoietic stem cells [13–15]. STAB-1 was implicated in lymphocyte adhesion and trafficking, angiogenesis and apoptotic cell clearance, therefore being crucial to maintain tissue homeostasis and resolving inflammation [16]. This SR has the ability to bind different ligands including modified low-density lipoproteins [17], phosphotidylserine expressed by apoptotic cells [18], secreted protein acidic and rich in cysteine (SPARC). Importantly, STAB-1 was previously found to bind Gram-positive and Gram-negative bacteria *in vitro* [19]. Furthermore, it is known that this receptor controls inflammatory activity, modulates T cell activation and also humoral immune response [20].

Here we address the role of STAB-1 in host defense against *Lm* infection and investigate the impact of STAB-1 deficiency on the host innate immune response against this bacterial pathogen. We reveal that STAB-1 KO mice display deregulated cytokine and chemokine expression, impaired recruitment of myeloid cells and increased susceptibility to *Lm* infection. In addition, whereas STAB-1 appears to promote bacterial uptake by macrophages, *Lm* infection induces the down regulation of STAB-1 expression and its delocalization from the host cell membrane.

## Materials and Methods

### Bacteria and cells

*Listeria monocytogenes* EGD (BUG 600) (*Lm*) and the non-pathogenic *Listeria innocua* (CLIP 11262) (*Li*) were grown in Brain Heart Infusion (BHI) (BD-Difco) at 37°C. *Lm* EGD transformed with pNF8-GFP plasmid (*Lm* EGD_GFP_) was grown in BHI supplemented with 5 μg/ml erythromycin. Human acute monocytic leukemia cells, THP-1 (ATCC TIB-202), were maintained in Roswell Park Memorial Institute (RPMI) 1640 medium (Lonza) supplemented with 10% foetal bovine serum (FBS) (BioWest). Before bacterial infection, THP-1 cells were differentiated with 10 nM phorbol 12-myristate 13-acetate for 48 h [21]. Murine macrophages J774 A.1 (ATTC TIB-67) and Raw 264.7 (ATTC TIB-71) were cultured in Dulbecco’s modified Eagle medium (DMEM) (Lonza), supplemented with 10% FBS. Human umbilical vein endothelial cells (HUVECs) were isolated and maintained in M199 culture medium supplemented with 10% FBS, heparin at 100 μg/ml and endothelial cell growth supplement (ECGS) at 30 μg/ml.

### Macrophage infection

Macrophages were incubated for 30 min with: 100 μg/ml of fucoidan (Sigma-Aldrich), 50 μg/ml of Poly(I) or Poly(C) (Santa-Cruz-Biotechnology). Cells were infected for 30 min with exponential-phase bacteria at a multiplicity of infection (MOI) of 2 and treated with 20 μg/ml of gentamicin (Lonza) for 60 min as described [22]. Raw macrophages were incubated with 5 μg/ml or 25 μg/ml of mouse-IgG (SC-2025) or anti-STAB-1 antibody (sc-98788) 1 h before bacterial infection at MOI of 50, during 30 min or 20 min plus 10 min with 50 μg/ml of gentamicin. Cells were washed and lysed for CFU quantification.

### Bone marrow-derived macrophages (BMDMs)

Mouse femurs were removed and flushed with Hank’s Buffered Salt Solution (HBSS-Lonza) as described [23]. Bone marrow cells were collected by centrifugation and cultured overnight in DMEM supplemented with 10 mM HEPES (Gibco), 1 mM sodium pyruvate (Lonza), 10% FBS and 10% L929 cell-conditioned medium (LCCM). Non-adherent cells were collected and seeded. Upon 4 days of differentiation, 10% of LCCM was added and on day 7 the medium was renewed. At day 10, latex beads of 1 μm (Invitrogen) (30 min of incubation) or exponential-phase bacteria at MOI 50 (20 min of infection plus 10 min with 50 μg/ml gentamicin) were added. Macrophages were washed and lysed for CFU quantification or used for immunofluorescence staining.

### RNA techniques

RNAs were extracted from non-infected and infected cells (TripleXtractor, GRISP), as recommended by the manufacturer. Purified RNAs were reverse-transcribed (iScript, Bio-Rad-Laboratories) and analysed by qPCR as described [24] or using specific primer probes (TaqMan). Gene expression data were analysed by comparative Ct method [25], normalized to *HPRT1* expression. For qualitative analysis, PCR was performed on cDNA (KAPA2G Mix, GRISP). Amplification products were resolved in 1% (w/v) agarose gel and analysed with GelDoc XR+ System (Bio-Rad Laboratories). Primers and probes are listed in Table S1.

### Immunofluorescence

*Lm* EGD_GFP_ infected BMDMs were fixed in 3% paraformaldehyde (15 min), quenched with 20 mM NH4Cl (1 h) and blocked with 1% BSA (sigma) in PBS (30 min). Cells were permeabilized with 0.1% Triton X-100 in PBS for 5 min and labelled with Alexa Fluor 647-conjugated phalloidin (Invitrogen), during 45 min on the dark. Cells were washed and slide preparations were mounted and dried at room temperature. Images were captured with an Olympus BX53 fluorescence microscope. The percentage of cells with intracellular bacteria or beads, and the number of intracellular bacteria or beads per cell were calculated. At least 300 cells were analysed for each sample in three independent experiments. Non-infected and *Lm*-infected HUVEC were incubated for 1 h with primary antibody rabbit anti-STAB-1 (1:100, Millipore), diluted in 0.2% saponin (Merck) supplemented with 1% BSA. Cells were washed in 0.2% saponin and incubated 45 min with secondary anti-rabbit Alexa 488 antibody (Invitrogen). DNA was counterstained with DAPI (Sigma) and actin labelled with TRITC-conjugated phalloidin. Images were collected with an Olympus BX53 fluorescence microscope and processed using ImageJ.

### Cell fractionation and Immunoblotting

Cytoplasmic and membrane fractions from non-infected and *Lm*-infected cells were obtained using the Subcellular Protein Fractionation kit (Thermo Scientific). Cell samples and homogenized spleens were diluted in Laemmli buffer, resolved by SDS-PAGE on 8% gels. Samples were transferred onto nitrocellulose membrane (Bio-Rad Laboratories), blocked and blotted with rabbit anti-STAB-1 (1:500, Millipore), followed by HRP-conjugated goat anti-rabbit IgG (1:2000, P.A.R.I.S). Signals were detected using ECL (Thermo-Scientific) and digitally acquired in a ChemiDoc XRS+ system (Bio-Rad Laboratories). Signal intensity was quantified using Image J.

### Cytokine ELISA

Lysis buffer 2x (200 mM Tris, 300 mM NaCl, 2% triton, pH 7.4) and Complete proteinase inhibitor (Roche) were added to homogenized organs for 30 min on ice. Supernatants were collected upon centrifugation and stored (−80°C). Mouse serum was recovered after blood centrifugation. Cytokine production was determined using murine ELISA kit (eBioscience).

### Flow cytometry

Mouse spleens were collected in ice-cold storage solution (PBS 2% FBS) and single-cell suspensions prepared using cell strainers (BD-Falcon). Cells were washed upon red blood cells lysis (150 mM NH_4_Cl, 10 mM KHCO_3_, pH 7.2 in H_2_O) and cell viability was assessed by trypan blue (Life-technologies) exclusion method. Peritoneal cells were collected by washing peritoneal cavities with 5 ml of storage solution, pelleted by centrifugation, washed and cell viability was assessed. Cells were labelled with brilliant violet 510-conjugated anti-CD11b, clone M1/70; BV 421-conjugated anti-CD11c, clone N418; allophycocyanin (APC)-conjugated anti-Ly6G, clone 1A8; APC with cyanin-7 (APC/Cy7)-conjugated anti-F4/80, clone BM8; and phycoerythrin (PE)-conjugated anti-Ly6C, clone HK1.4 (BioLegend). Data were acquired in a FACS Canto II flow cytometer (BD-Biosciences) and analysed using FlowJo software (TreeStar Inc.). To determine cell numbers, event number for each cell population was normalized to the total cell number.

### Animal infections

STABILIN-1 full knock-out (STAB-1 KO) mice and their wild-type (WT) littermates, both with a C57BL/6N, 129SvJ mixed background have been described [26]. Infections were done as described [27]. Briefly, intravenous infections were performed through the tail vein with 5×10^5^ colony-forming units (CFUs) in PBS. Mice were euthanized 72h post-infection, spleens and livers were aseptically collected and CFUs counted. Blood was recovered from mice heart. Mouse survival was assessed upon intravenous infection of 10^5^ CFUs. Animals were intraperitoneally injected with 10^5^ CFUs (*Lm*) or 5 mg/kg of LTA from *Staphylococcus aureus* (L2515 Sigma) in PBS and euthanized 6h or 24h later. Animal procedures followed European Commission (directive 2010/63/EU) and Portuguese (Decreto-Lei 113/2013) guidelines and were approved by the IBMC Ethics Committee and Direção Geral de Veterinária (license 015301).

### Statistics

Statistics were carried out with Prism (GraphPad), using unpaired two-tailed Student’s *t*-test to compare means of two groups, and one-way ANOVA with Tukey’s post-hoc test for pairwise comparison of means from more than two groups, or with Dunnett’s post-hoc test for comparison of means relative to the mean of a control group.

## Results

### Scavenger Receptors are required for Lm uptake by macrophages

To evaluate the overall role of SRs in *Lm* uptake by eukaryotic cells, we chemically saturated SRs using different pleiotropic compounds (fucoidan, Poly (I)) known to inhibit SRs [28], before *Lm* infection of human (THP-1) macrophages. Pre-treatment of THP-1 cells with fucoidan severely impaired *Lm* uptake when compared to non-treated cells (Figure 1A). In addition, the number of intracellular bacteria was also reduced upon SR saturation with Poly (I), but not with its corresponding negative control Poly (C) (Figure 1A). In agreement, pre-treatment of murine macrophage-like cells (Raw and J774 cell lines) with fucoidan also compromised *Lm* uptake (Figure 1B). These data suggested a role for SRs in *Lm* uptake by macrophages.

**Figure 1.**
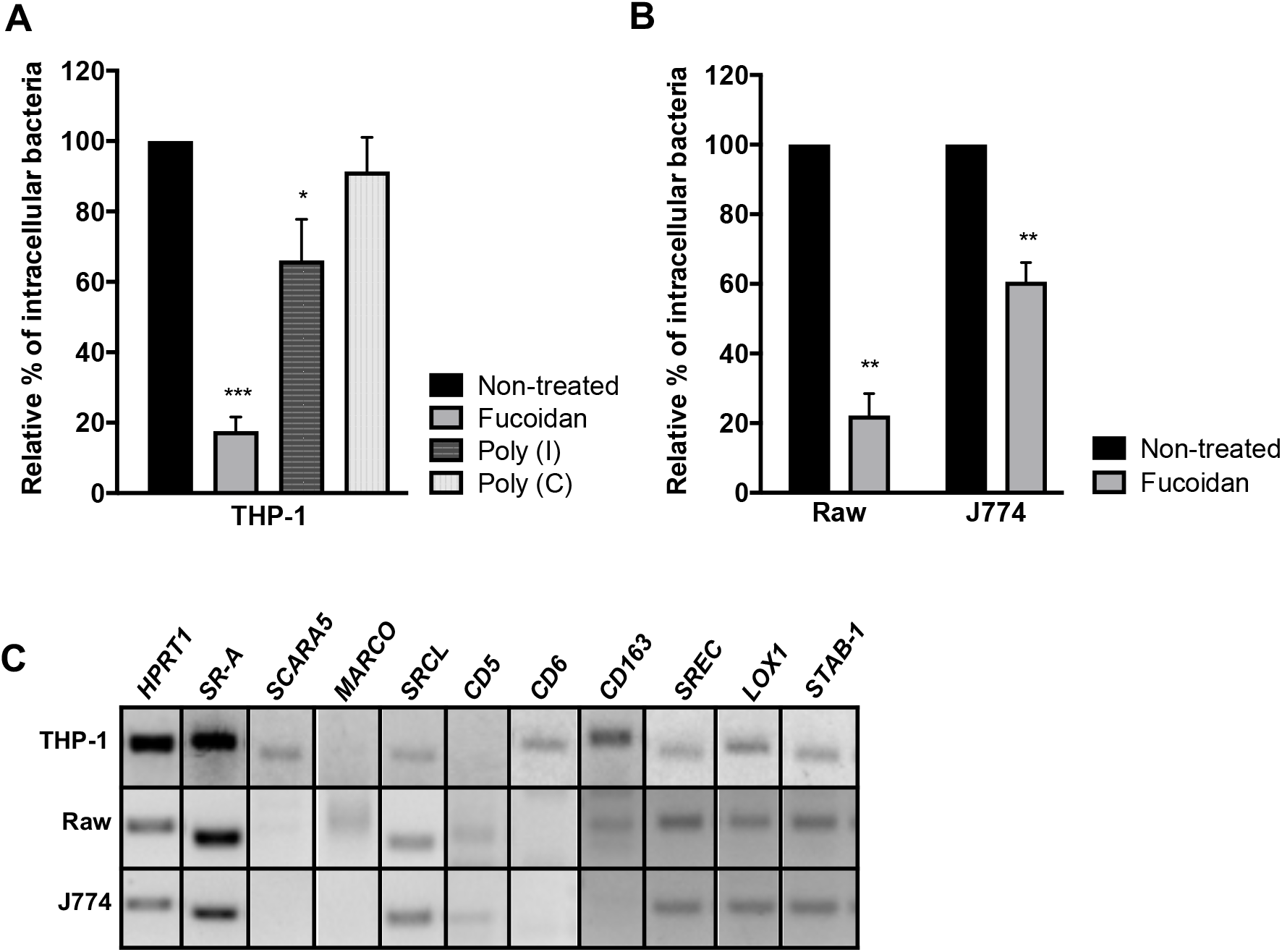
SRs are required for *Lm* uptake by macrophages. (A-B) Chemical saturation of SRs impairs bacterial uptake by macrophages-like cells. (A) Human THP-1, (B) murine Raw and J774 macrophage-like cells were left untreated or pre-treated with fucoidan, Poly(I) and its control Poly(C), infected by *Lm* for 30 min, incubated with gentamicin, washed and lysed to quantify intracellular bacteria. Values are expressed relative to values in non-treated cells, arbitrarily fixed to 100%. Values are mean ± SD of three independent assays. \**p*<0.05; \*\**p*<0.01; \*\*\**p*<0.001. (C) SR gene expression was assessed by RT-PCR analysis on THP-1, Raw and J774 total RNAs, using *HPRT1* as reference gene. Band panel for each cell line is representative of two assays.

To identify SRs potentially involved in *Lm* uptake by macrophages, we assessed SR expression profiles by analysing total RNAs isolated from human and murine macrophage cell lines. Our analysis revealed that, although some of the selected SRs appeared broadly expressed, each cell line presented a specific SR expression profile (Figure 1C). In the tested conditions, *SR-A*, *SRCL*, *SREC*, *LOX1* and *STAB-1* appeared to be expressed in all cell lines. SR-A was previously proposed to play a crucial role in host defense against *Lm* infection [9]. Interestingly, STAB-1 was previously implicated in lymphocyte transmigration and apoptotic cell clearance [16], and shown to bind Gram-positive and Gram-negative bacteria *in vitro* [19]. Since the involvement of STAB-1 in infectious processes was never assessed so far, we further explore its potential role on *Lm* infection.

### STAB-1 is required for Lm uptake by macrophages

During infection, systemic bacteria are sequestered by phagocytes both in the liver and spleen [29]. SRs are expressed by macrophages and may function as phagocytic receptors for bacteria [19]. Aiming at understanding the role of STAB-1 in *Lm* uptake, we pre-incubated Raw macrophages with anti-IgG (negative control) or anti-STAB-1 antibody before *Lm* infection. While the percentage of adherent bacteria was similar between IgG- and anti-STAB-1 treated cells, the percentage of intracellular *Lm* diminished upon macrophage treatment with anti-STAB-1 antibody (Figure 2A). These data suggest that saturating STAB-1 on the surface of macrophages reduces *Lm* uptake. To further address the role of STAB-1 in *Lm* uptake by macrophages, bone marrow-derived macrophages (BMDMs) from both WT and STAB-1 KO mice were infected with *Lm*. As compared to WT, STAB-1 KO macrophages displayed decreased numbers of intracellular *Lm* (Figure 2B). Immunofluorescence quantifications of the percentage of infected cells and the number of intracellular bacteria per cell confirmed the reduced capacity of STAB-1 KO macrophages to uptake *Lm* (Figure 2C). STAB-1 KO macrophages also displayed a slight phagocytosis defect of non-pathogenic *Listeria* (*Listeria innocua* - *Li*) [30], as well as of latex beads (Figure 2D). However this defect appeared more pronounced for *Lm* than for *Li* or beads. STAB-1 appears thus be involved in the uptake of foreign bodies by macrophages.

**Figure 2.**
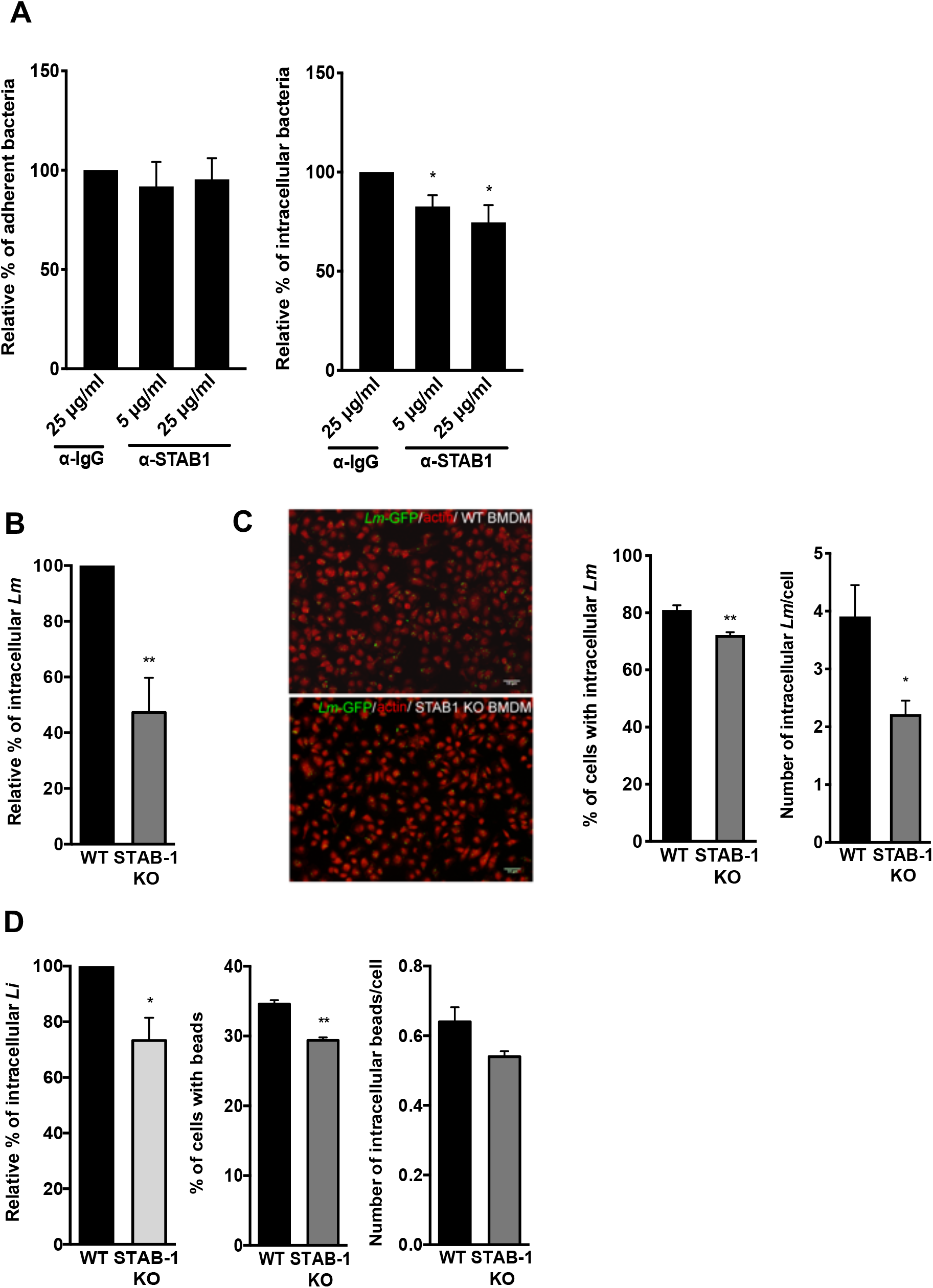
STAB-1 is required for *Lm* uptake by macrophages. (A) Impact of STAB-1 on *Lm* adhesion and entry into macrophages. Raw were pre-treated with an anti-IgG (SC-2025) or anti-STAB-1 antibody, at 5 μg/ml and 25 μg/ml (sc-98788) before *Lm* infection. Adherent and intracellular bacteria were quantified. Values are expressed relative to values in IgG-treated cells, arbitrarily fixed to 100%. (B) Quantification of intracellular bacteria in WT and STAB-1 KO BMDMs infected with *Lm* for 30 min. Values are expressed relative to WT arbitrarily fixed to 100%. (C) Immunofluorescence images of WT and STAB-1 KO BMDMs infected with *Lm*-GFP (green) for 30 min. Actin is labelled with TRITC-conjugated phalloidin. Scale bar, 10 μm. Quantification of the percentage of cells with intracellular *Lm* and of the number of intracellular *Lm* in BMDMs upon 30 min of infection. Values are mean ± SD of three to four independent experiments. Statistical significance is indicated as compared to WT BMDMs. \**p*<0.05; \*\**p*<0.01. (D) Quantification of intracellular *Li,* the percentage of cells with beads and the number of intracellular beads per cell in WT and STAB-1 KO BMDMs infected for 30 min. Values are mean ± SD of three independent experiments. Statistical significance is indicated as compared to WT BMDMs. \**p*<0.05; \*\**p*<0.01

### STAB-1 has a protective role against Lm infection

To investigate the contribution of STAB-1 during *Lm* systemic infection *in vivo*, WT and STAB-1 KO mice were intravenously infected with *Lm*. Three days later, mice were euthanized and bacterial loads in spleens and livers were quantified. Bacterial numbers appeared significantly higher in the organs of STAB-1 KO mice (Figure 3A), demonstrating a role for STAB-1 in the control of *Lm* infection. To test if this defect in the control of infection may lead to increased mortality, mice were intravenously infected with a lower dose of *Lm* and survival was monitored over time. Whereas WT mice survived throughout the infection, mortality in STAB-1 KO mice reached 80% by day 18 (Figure 3B).

**Figure 3.**
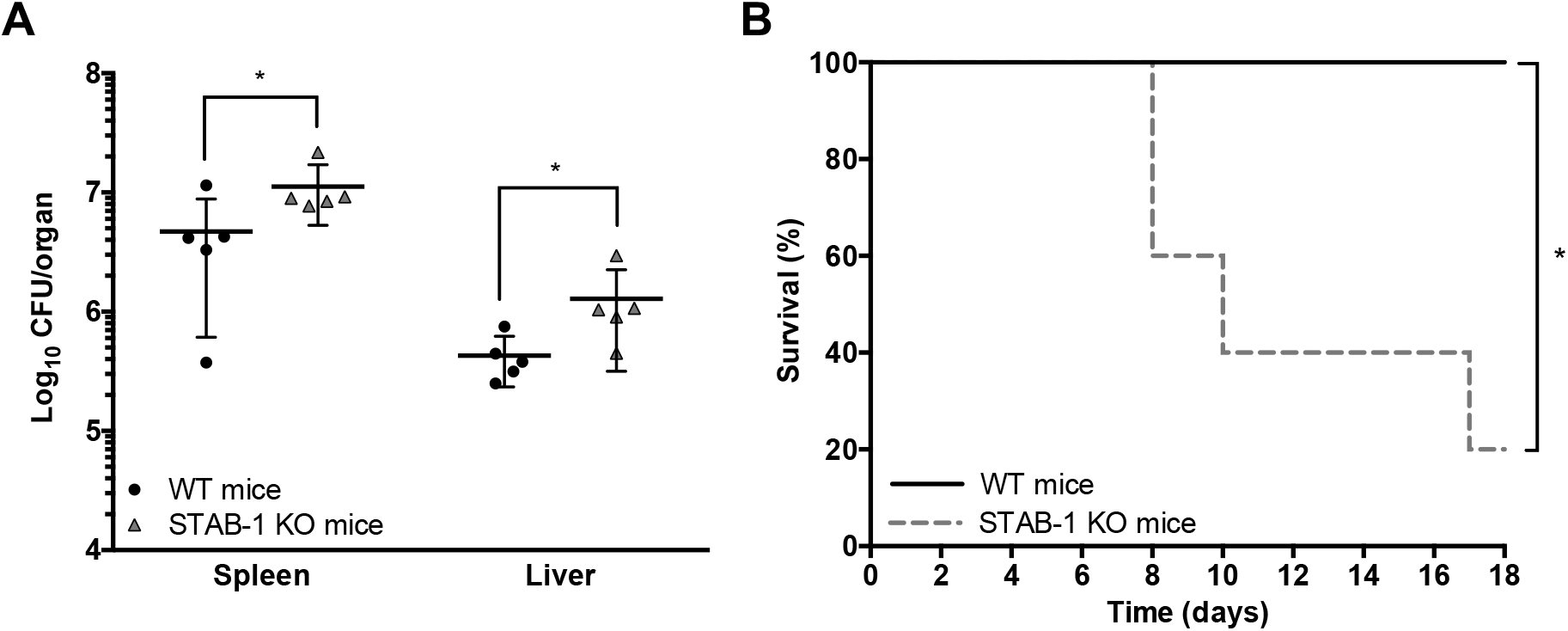
STAB-1 have a protective role against *Lm* infection. (A) Quantification of viable bacteria in spleens and livers recovered from WT and STAB-1 KO mice, three days after intravenous infection of 5×10^5^ CFU of *Lm.* Data are presented as scatter plots, each animal is represented by a dot and the mean is indicated by a horizontal line. (B) WT and STAB-1 KO mice survival after intravenous inoculation of 10^5^ CFU of *Lm* (n=5). \**p*<0.05.

Altogether, our data indicate that STAB-1 promotes protection against *Lm* infection.

### STAB-1 is required for an efficient inflammatory response and immune cell accumulation in Lm-infected spleens

Mouse infection by *Lm* induces a robust innate inflammatory response that restricts bacterial growth prior to the development of protective T cell responses. Early protective immunity against *Lm* relies on the production and balance of pro-inflammatory cytokines, such as TNF-α and IL-6, and anti-inflammatory cytokines, such as IL-10 [31]. To analyse the potential role for STAB-1 in the production of microbicidal mediators in response to *Lm* infection, WT and STAB-1 KO mice were intravenously infected with *Lm*. The production of cytokines in the spleens, livers and sera of *Lm*-infected mice was evaluated by ELISA three days post-infection. As compared to WT animals, infected STAB-1 KO mice produced lower levels of TNF-α, IL-6 and IL-10 (Figure 4A). Importantly, this reduction of the cytokine levels between WT and STAB-1 KO mice is not observed in absence of infection (Figure S1). These results indicate that STAB-1 plays a role in the coordinated cytokine production elicited by *Lm* infection in targeted mouse organs.

**Figure 4.**
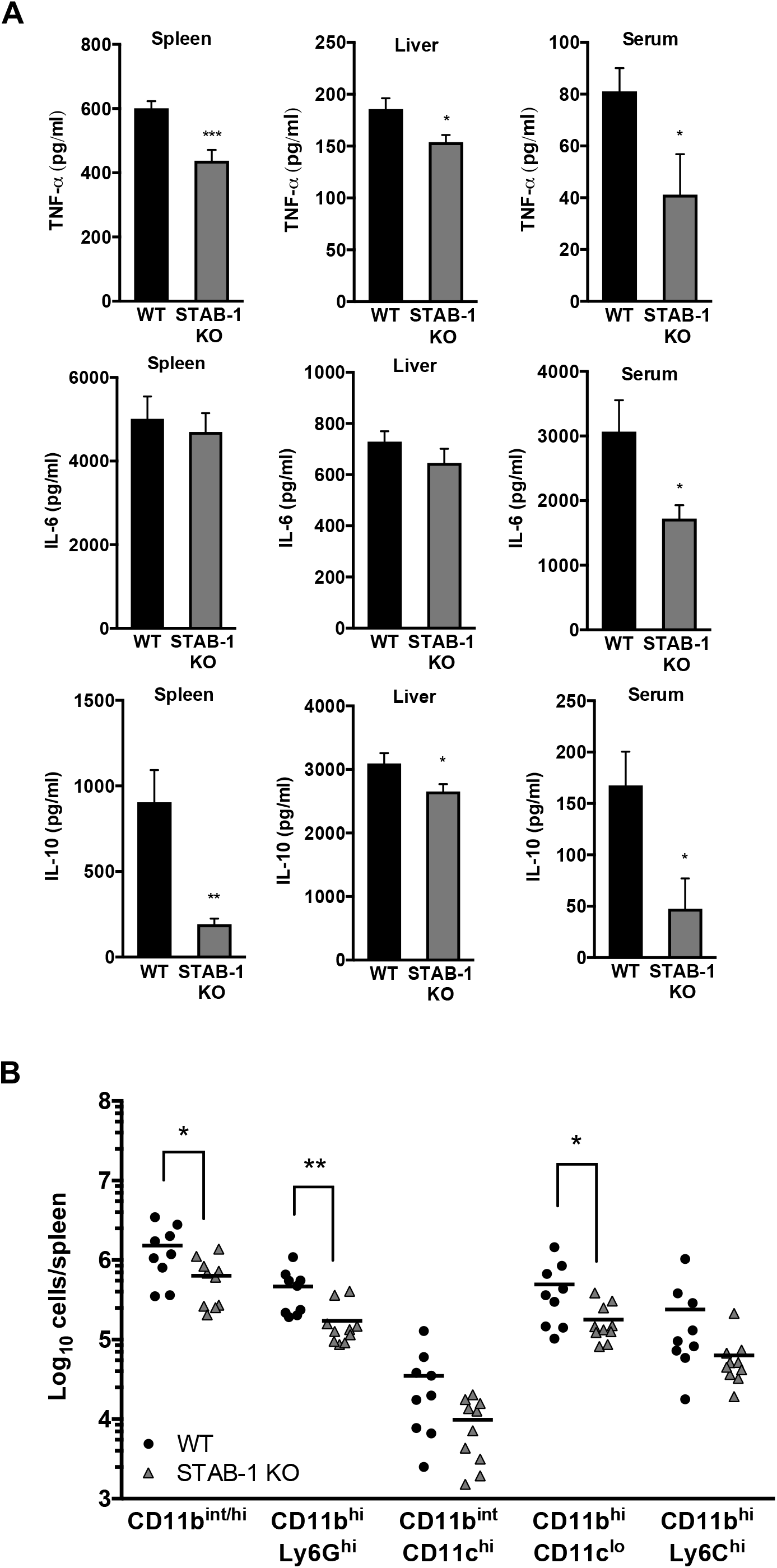
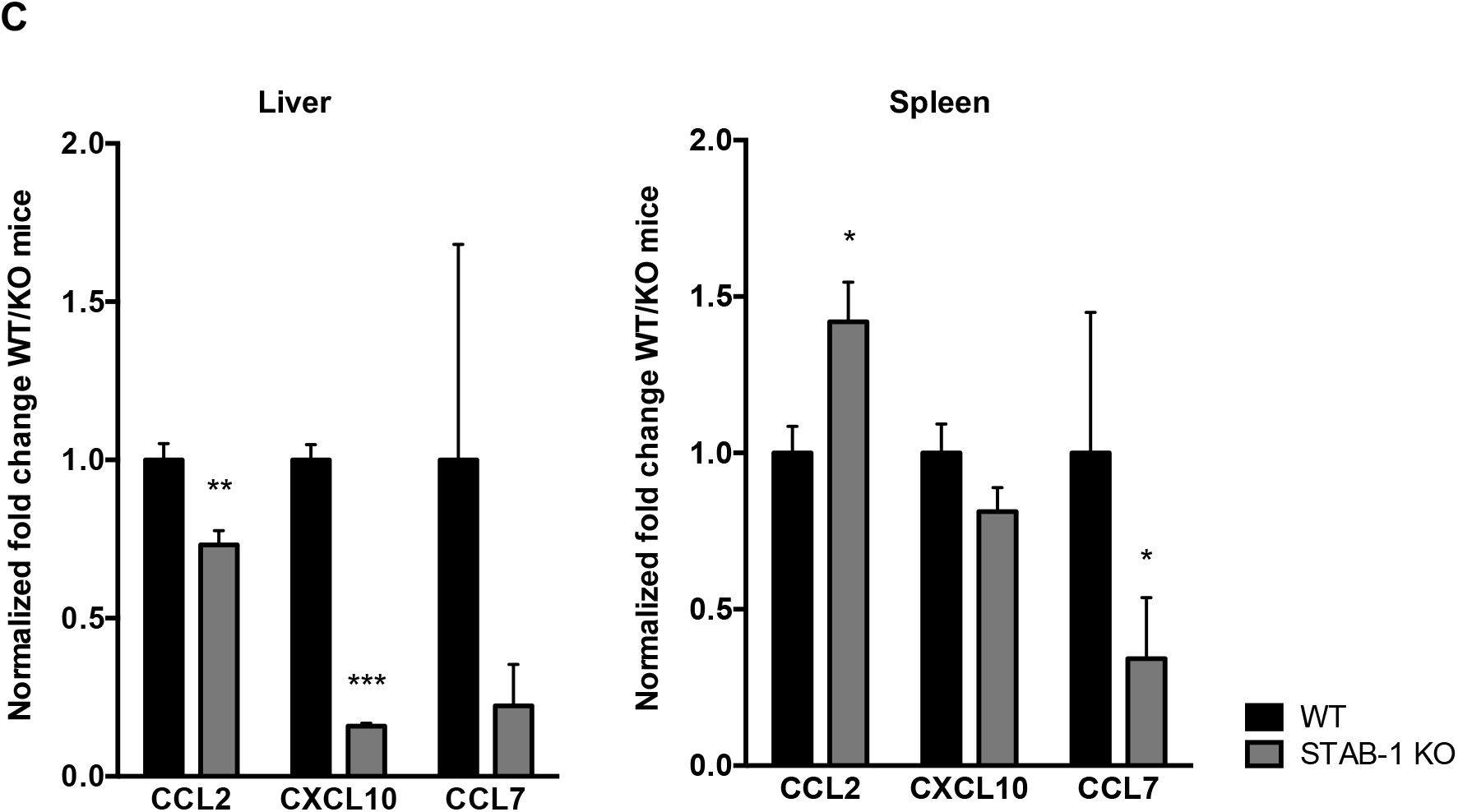
STAB-1 is required for an efficient inflammatory response and immune cell accumulation in *Lm*-infected spleens. (A) WT and STAB-1 KO mice were intravenously infected with 5×10^5^ CFU of *Lm.* Mice were sacrificed at day three post-infection and spleen, liver and serum were collected. Levels of TNF-α, IL-6 and IL-10 were measured by ELISA. Data are represented as an average of ten mice from two independent experiments per group. \**p*<0.05; \*\**p*<0.01; \*\*\**p*<0.001. (B) Spleen cells from *Lm* (5×10^5^ CFU) infected WT and STAB-1 KO mice were isolated and analysed by flow cytometry. Total numbers of myeloid cells (CD11b^int/hi^), neutrophils (CD11b^hi^Ly6G^hi^), dendritic cells (CD11b^int^CD11c^hi^), macrophages (CD11b^hi^CD11c^lo^) and inflammatory macrophages (CD11b^hi^Ly6C^hi^) are shown. Data are presented as scatter plots, with each animal represented by a dot and the mean indicated by a horizontal line. \**p*<0.05; \*\**p*<0.01. (C) WT and STAB-1 KO mice were intravenously infected with 5×10^5^ CFU of *Lm.* Mice were sacrificed at day three post-infection to recover spleens and livers. The expression of chemokines CCL2, CXCL10 and CCL7 was quantified by qRT-PCR. Data are represented as an average of ten mice from two independent experiments per group. \**p*<0.05; \*\**p*<0.01; \*\*\**p*<0.001.

*Lm* entering the bloodstream are rapidly taken up by various myeloid cells in tissues. In the spleen, bacteria are filtered by resident myeloid cells, including dendritic cells and professional phagocytes [32]. Inflammatory stimuli also induce the recruitment of inflammatory macrophages to infected tissues [33]. As STAB-1 appears to regulate the production of inflammatory cytokines in response to *Lm* infection, in particular in the spleen, we hypothesized that STAB-1 could impact innate immune cells recruitment to the infected spleen, a major site of bacteria replication. To test this hypothesis, WT and STAB-1 KO mice were intravenously infected with *Lm*, and three days post-infection, single-cell spleen suspensions were analysed regarding myeloid cell populations by flow cytometry. As compared to WT infected mice, *Lm*-infected STAB-1 KO mice showed a clear defect on myeloid CD11b^int/hi^ cells, which resulted from the diminished number of neutrophils (CD11b^hi^Ly6G^hi^) and macrophages (CD11b^hi^CD11c^lo^) (Figure 4B). Interestingly, within the macrophage population, the number of inflammatory macrophages (CD11b^hi^Ly6C^hi^) was also reduced in infected STAB-1 KO animals (Figure 4B). In absence of infection, spleens of WT and STAB-1 KO mice showed comparable myeloid cell populations (Figure S2). Taken together, these data show that STAB-1 is important in controlling the recruitment of neutrophils and macrophages to the spleen of *Lm*-infected mice.

The migration and positioning of immune cells in tissues in response to infection is mainly controlled by chemokines [34]. We thus analysed the expression of neutrophil- and monocyte-attracting chemokines in *Lm*-infected murine organs. In infected STAB-1 KO mouse livers, the expression of all chemokines tested was decreased as compared to WT infected mice (Figure 4C, left graph). In infected spleens, the expression of CCL7 and CXCL10 was also decreased in STAB-1 KO mouse spleens, whereas the expression of CCL2 was increased as compared to WT (Figure 4C, right graph). Differences observed between organs might be the result of niche/microenvironment disparities. Altogether, these results indicate a role for STAB-1 in the recruitment of immune cells to *Lm*-infection sites possibly through the expression control of attracting chemokines. These data also corroborate the role of CCL7 and CXCL10 in the recruitment of inflammatory monocytes to the spleen.

### STAB-1 is important for early myeloid cells recruitment in response to Lm infection

The early recruitment of immune cells to infected tissues was shown to be crucial for an effective innate immune response against *Lm* [33]. To evaluate the role of STAB-1 in the early trafficking of myeloid cells to the site of *Lm* infection, WT and STAB-1 KO mice were intraperitoneally infected with *Lm*. Exudate cells from the peritoneal cavity were recovered 6h or 24h post-infection to evaluate myeloid cell populations by flow cytometry. When compared to non-infected mice, *Lm* infection appeared to trigger the recruitment of cells to the focus of infection, mainly neutrophils (CD11b^hi^Ly6G^hi^) and inflammatory macrophages (CD11b^hi^Ly6C^hi^) (Figures 5A-C). While similar cell populations were detected in non-infected WT and STAB-1 KO mice (Figure 5A), we observed a defect recruitment of myeloid cells in STAB-1 KO when compared to WT mice, both at 6h and 24h after *Lm* infection (Figures 5B-C).

**Figure 5.**
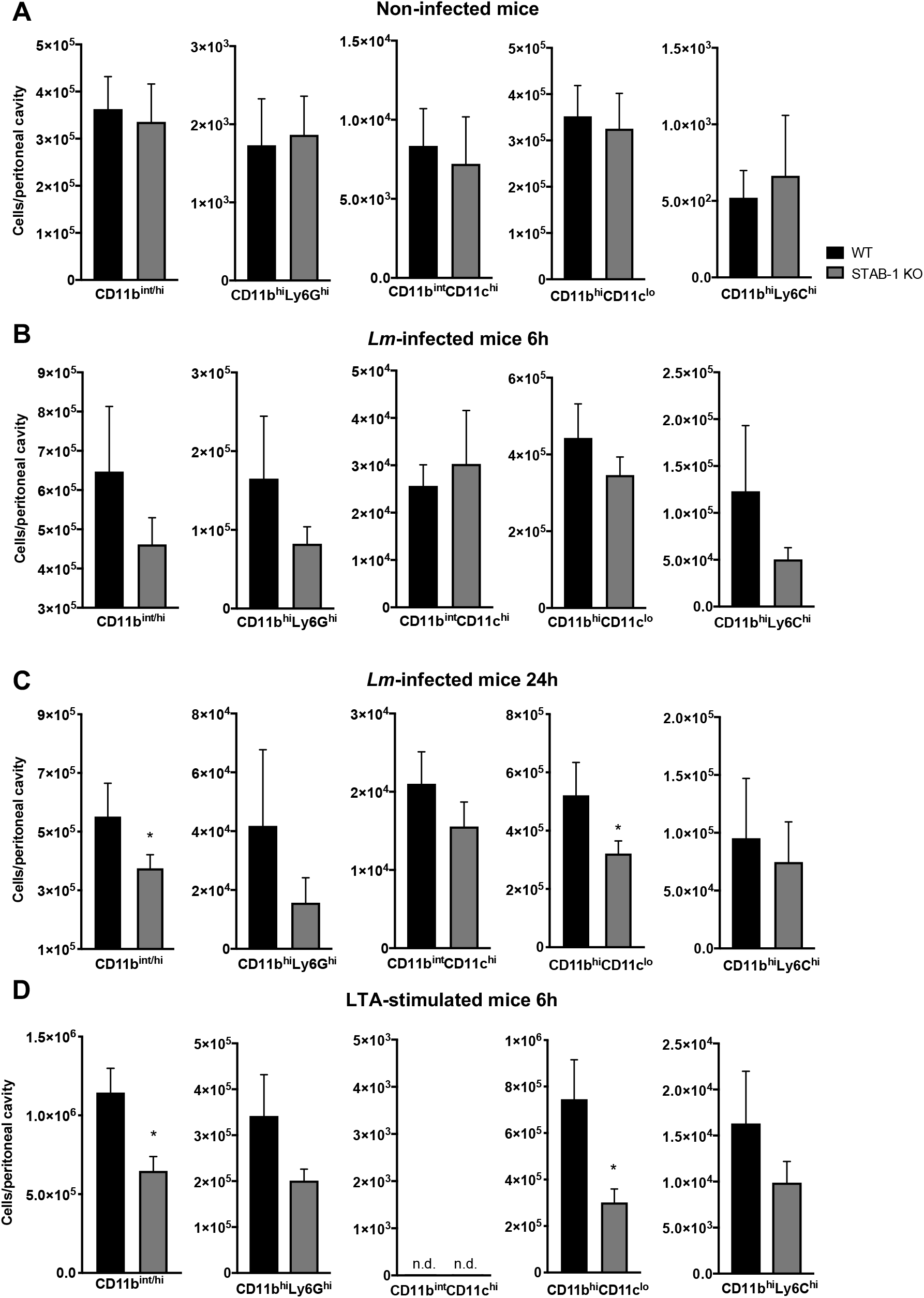
STAB-1 is important for early myeloid cells recruitment in response to *Lm* infection. (A-D) Single-cell suspensions recovered from the peritoneal cavity of WT and STAB-1 KO mice were analysed by flow cytometry to evaluate cell populations. (A) Non-infected animals. (B-C) Mice intraperitoneally infected with 10^5^ CFU of *Lm* for (B) 6 h or (C) 24 h. (D) Mice intraperitoneally injected with purified LTA (5 mg/ml) for 6 h. Data are represented as an average of two independent experiments, with at least six mice per group. \**p*<0.05; \*\*\**p*<0.001

Lipoteichoic acids are components of Gram-positive bacteria and potent inducers of inflammation. They stimulate immune cells and induce the migration of myeloid cells to the mouse abdominal cavity when injected intraperitoneally [35, 36]. We used this experimental model to confirm the involvement of STAB-1 in the recruitment of innate immune cells to the infection site. Purified LTA were intraperitoneally injected into WT and STAB-1 KO mice and, 6h post-stimulation, exudate cells from the peritoneal cavity were recovered to evaluate myeloid cell populations. In response to LTA, STAB-1 KO mice showed a significant reduction in the myeloid cell (CD11b^int/hi^) population when compared to WT mice, which correlates to a decreased recruitment of neutrophils (CD11b^hi^Ly6G^hi^), macrophages (CD11b^hi^CD11c^lo^) and inflammatory macrophages (CD11b^hi^Ly6C^hi^) (Figure 5D). Altogether, these results indicate that, *in vivo,* STAB-1 potentiates the recruitment of immune cells to the infection site upon an inflammatory stimulus.

### STAB-1 expression is decreased and re-localized in response to Lm infection

Since STAB-1 appeared to restrain *Lm* infection by regulating cytokine and chemokine production and controlling myeloid cell recruitment, we investigated the potential impact of *Lm* infection on STAB-1 expression. We analysed *STAB-1* expression in murine macrophage-like cells (J774) in response to *Lm* infection and showed a slight decrease of *STAB-1* expression in infected as compared to non-infected macrophages (Figure 6A). As observed in macrophage cell lines, *Lm* infection also induced the down-regulation of STAB-1 expression in mouse BMDMs at the RNA and protein level (Figure 6A and 6B). Interestingly, this down-regulation of STAB-1 expression in BMDMs was only observed with pathogenic *Listeria* (*Lm*) and not with the non-pathogenic species (*Li*) (Figure 6A and 6B).

**Figure 6.**
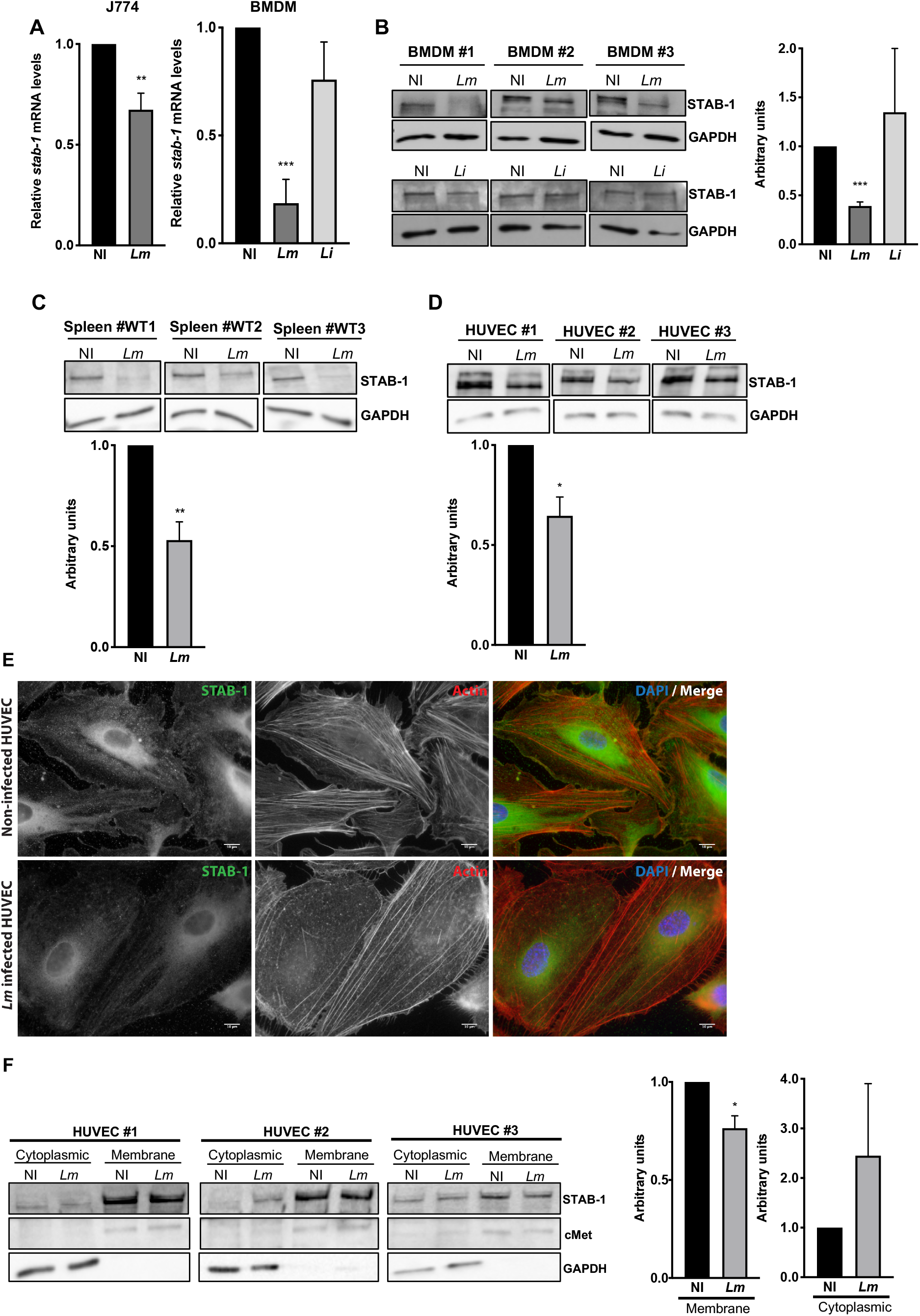
*STAB-1* expression is decreased and re-localized in response to *Lm* infection. (A) Assessment of STAB-1 expression by (A) quantitative RT-PCR and (B) Western Blot. (A) Quantification of *STAB-1* mRNA levels on RNAs extracted from J774 and BMDMs infected with *Lm* for 30 min. *STAB-1* expression levels in infected conditions were normalized to those in non-infected BMDMs, arbitrarily fixed to 1. (B-D) Independent immunoblots to detect STAB-1 protein in (B) BMDMs left uninfected (NI) or infected with *Lm* or *Li* for 30 min, (C) spleen of NI and *Lm*-infected mice for three days (5×10^5^ CFU), (D) NI and *Lm*-infected HUVECs. Immunoblots quantification of STAB-1 signal intensity in NI and infected conditions, normalized to GAPDH. (E) Immunofluorescence images of NI and *Lm* infected HUVECs, stained with an anti-STAB-1 antibody. DNA counterstained with DAPI and actin labelled with TRITC-conjugated phalloidin. Scale bar, 10 μm. (F) Immunoblots to detect STAB-1 protein in the cytoplasmic and membrane fractions of NI and *Lm*-infected HUVECs upon cell fractioning. cMet was used as a membrane loading protein control and GAPDH as a cytoplasmic loading protein control. Immunoblots quantification of STAB-1 signal intensity in NI and infected conditions, normalized to cMet or GAPDH.

As we showed that STAB-1 is required for an efficient immune response during *Lm* infection of mouse spleens, we thus assessed the impact of *Lm* infection on STAB-1 expression *in vivo,* in splenic tissue. The analysis indicated that *Lm* infection also induced a large decrease of STAB-1 expression in infected mouse tissues (Figure 6C). In the spleen, STAB-1 is not expressed by splenic macrophages but mainly by endothelial cells [13] that were shown to be active participants in the inflammatory response during *Lm* infection [37]. Therefore, we evaluated whether *Lm* infection could impact STAB-1 expression on endothelial cells (HUVECs) and showed again a significant decrease of STAB-1 levels in infected cells (Figure 6D). Immunofluorescence analysis corroborated the down-regulation of STAB-1 expression upon *Lm* infection of endothelial cells (Figure 6E). In addition, by microscopy, we also observed a de-localization of STAB-1 from the host cell membrane upon infection (Figure 6E). This was further confirmed by cell fractionation that showed a significant reduction of membrane STAB-1 in *Lm*-infected HUVECs when compared to non-infected cells (Figure 6F).

Together, these results indicate that infection by pathogenic *Listeria* induces a down-regulation of STAB-1 expression in infected cells and tissues. This down-regulation is accompanied by a de-localization of STAB-1 from the host cell membrane.

## Discussion

Scavenger receptors (SRs) are transmembrane cell surface glycoproteins restricted to macrophages, dendritic cells, endothelial cells and a few other cell types [38]. Whereas SRs were initially defined by their ability to bind modified low-density lipoproteins, several SRs were since demonstrated to play an important role in innate immune defenses [39].

Here, we show for the first time the important role of the SR STAB-1 in the host protection against bacterial infection. We demonstrate that, during an infection by *Listeria monocytogenes* (*Lm*), STAB-1 is not only required for bacterial uptake by macrophages, but also for an efficient inflammatory response, immune cell accumulation, and early myeloid cells recruitment to the infection site. Interestingly, we also show that infection by pathogenic *Listeria* induces the down-regulation of STAB-1 expression and its de-localization from the cell membrane, suggesting a bacterial active virulence process targeting STAB-1 aiming to promote infection. STAB-1 appears thus as a new important player in the host protection against a major Gram-positive food-borne pathogen.

SRs were previously shown to represent an important part of the innate immune defense, in particular by acting as phagocytic receptors for microorganisms [40–42]. In addition, STAB-1 was previously found to bind Gram-positive and Gram-negative bioparticles *in vitro* [19], and was described to be a phagocytic receptor mediating efferocytosis by recognizing phosphatidylserine on apoptotic cells [43]. We show here the reduced ability of STAB-1 KO macrophages to uptake not only *Lm*, but also non-pathogenic bacteria and beads. This could suggest a role for STAB-1 in the general phagocytic process. However, it was previously documented that antibody blockade or absence of STAB-1 is sufficient to skew macrophages from an anti-inflammatory to a more pro-inflammatory phenotype [17], these later being inherently less phagocytic for *Lm* or latex beads than their anti-inflammatory counterparts [44]. This could be responsible, at least in part, for the decreased phagocytic capacity observed for STAB-1 KO macrophages.

We report here that STAB-1 contributes in the host response against *Lm* infection by controlling cytokine and chemokine production, thus controlling myeloid cell recruitment. SRs are strong players in the regulation of inflammation, such is the case of SR-A in *Neisseria meningitidis* and *Porphyromonas gingivalis* infections [45, 46] or CD36 in response to *Staphylococcus aureus* [47]. STAB-1 was previously shown to control the activation of several pro-inflammatory cytokines in human monocytes [48]. Here, we show that infected STAB-1 KO mice produced reduced serum, liver and splenic levels of IL-6 and TNF-α as compared to WT mice, suggesting that STAB-1 participates in the regulation of the inflammatory cytokine response in *Lm-*targeted mouse organs upon infection. In agreement, IL-6- and TNF-α-deficient mice were shown to be more susceptible to *Lm* infection, with increased bacterial burden in the spleen and liver, and deficient neutrophil recruitment into the blood [49, 50]. Surprisingly, we also observed an IL-10 decreased expression in STAB-1 KO mice upon *Lm* infection. IL-10 is a potent inhibitor of innate immunity and IL-10 deficiency was shown to improve resistance to *Lm* infection [51].

We show here that STAB-1 appears as an important regulator of the pro-/anti-inflammatory cytokine balance in response to *Lm* infection. *Lm* uses a balance of pro- and anti-inflammatory mechanisms to promote infection while inducing little inflammation in the host, both at the intestinal level and systemically [52]. In a rodent model of intestinal infection, differential induction of pro- or anti-inflammatory responses depending on the cell type used for entry were observed [53]. Thus a picture emerges that the modulation of pro- and anti-inflammatory properties at the cellular level has important consequences for the course of infection when monitored in the complex host environment. Whereas STAB-1 was previously proposed as an immunosuppressive molecule, suggesting that STAB-1 may dampen pro-inflammatory reactions *in vivo* [48], our results rather tend to indicate a pro-inflammatory role for STAB-1 during *Lm* infection. In the context of sepsis, where the pro-inflammatory response predominates, STAB-1 was previously proposed to be both an immunosuppressive player to down-regulate hyper-inflammation at early stages and to act as a vascular barrier keeper in later stages of sepsis. STAB-1 could thus appear as a regulator of inflammatory processes, acting both as a pro- and anti-inflammatory molecule depending of the context and localization in the host.

Our findings also indicate that STAB-1 plays a role in the recruitment of myeloid cells in infected organs, a recruitment that appears to be dependent on chemokine expression, in particular CXCL10 and CCL7. CXCL10 was previously involved in immune cell migration, differentiation and activation [54], and is induced by TNF-α [55]. The reduced levels of CXCL10 in STAB-1 KO mice upon *Lm* infection might thus correlate with the concomitant decreased expression of TNF-α. Interestingly, CXCL10 has been shown to have direct antibacterial properties similar to α-defensins, in particular against *Lm* [56]. During *Lm* infection, the recruitment of inflammatory macrophages from bone marrow to sites of microbial infection was shown to be dependent on CCR2, a chemokine receptor that responds to CCL2 and CCL7 [57]. In our experimental model, CCL7 seems to play a more prominent role in the STAB-1-dependent recruitment of myeloid cells to *Lm* infected organs. However, the slight increase of CCL2 expression observed in the spleen of *Lm*-infected STAB-1 KO mice could also suggest a role of STAB-1 in myeloid cell chemotaxis.

We show that the deficiency on pro-inflammatory cytokine production and the defect on myeloid cell recruitment, which are crucial for the initial control of bacterial replication, lead to higher bacterial loads in the spleen and liver of STAB-1 KO mice. Neutrophils and macrophages, which are effective microbicidal cells, are among the first cells involved in the *Lm*-immune response. Mice deficient for these cells present increased bacterial burden and mortality [58]. The reduced capacity of STAB-1 KO mice to fight *Lm* infection appears thus to be more related to a deficiency in the recruitment of myeloid cells to target organs, than to a killing deficiency. In agreement, STAB-1, which was shown to be absent from all splenic macrophages, including red pulp, marginal zone and metallophilic macrophages, is solely expressed by the vascular endothelium [59], and is known to be involved in the transmigration of immune cells [14, 15]. Nevertheless, impaired control of the infection by STAB-1 KO mice may not be only due to reduced myeloid cell recruitment to the sites of infection. STAB-1 could be required for the proper migration, position and function of other immune cells, in particular CD8α^+^ dendritic cells of the splenic marginal zone that were shown to be an obligate cellular entry point for a productive infection by *Lm* [60]. However, it was previously shown that the frequencies of splenic CD4+ and CD8+ T cells were comparable between wildtype and STAB-1 KO mice [20]. Our work focuses on the importance of STAB-1 in the early immune response of *Lm*, but further studies need to be performed to understand weather the deficiency of STAB-1 in the context of *Lm* infection interferes with T cell priming and the generation of T cell memory.

Whereas STAB-1 appears as an important player in the host protection against *Lm*, we also show that infection by this bacterial pathogen induces a decreased expression of STAB-1 in macrophages and endothelial cells but also *in vivo* in infected mice spleen, which is a major target organ for *Lm* replication. The expression of some other SRs, such as SR-A, MARCO and LOX-1, was also reported to be modulated by microbial infection, either favoring host immune response or promoting pathogen survival [45, 61, 62]. In particular, a marked expression of MARCO was observed in response to *Leishmania major* and *Lm* infections [10, 63]. Interestingly, we also found that *Lm* infection leads to a delocalization of STAB-1 from the membrane of endothelial cells. Importantly, the down-regulation of STAB-1 expression was not observed with the nonpathogenic specie *Listeria innocua*, that essentially differs from *Lm* by the absence of major virulence factors [30]. This suggests an active mechanism driven by *Lm* virulence factors to control STAB-1 expression/localization, diminish host protective responses and promote infection. However, the identification of the specific virulence factors potentially involved in this process requires further investigation.

Here, we highlight for the first time that STAB-1 plays a protective role during *Lm* infection. By regulating the inflammatory response and the recruitment of myeloid cells, STAB-1 appears as a new SR with an important role for the host response against *Lm* infection. Amplifying STAB-1-mediated host defenses may represent an innovative strategy against Gram-positive pathogens.

## Acknowledgements

We thank Rui Appelberg for PhD co-supervision of R.P. and J.P., for helpful discussions and for critical reading of this manuscript.

---

**Figure S1.**
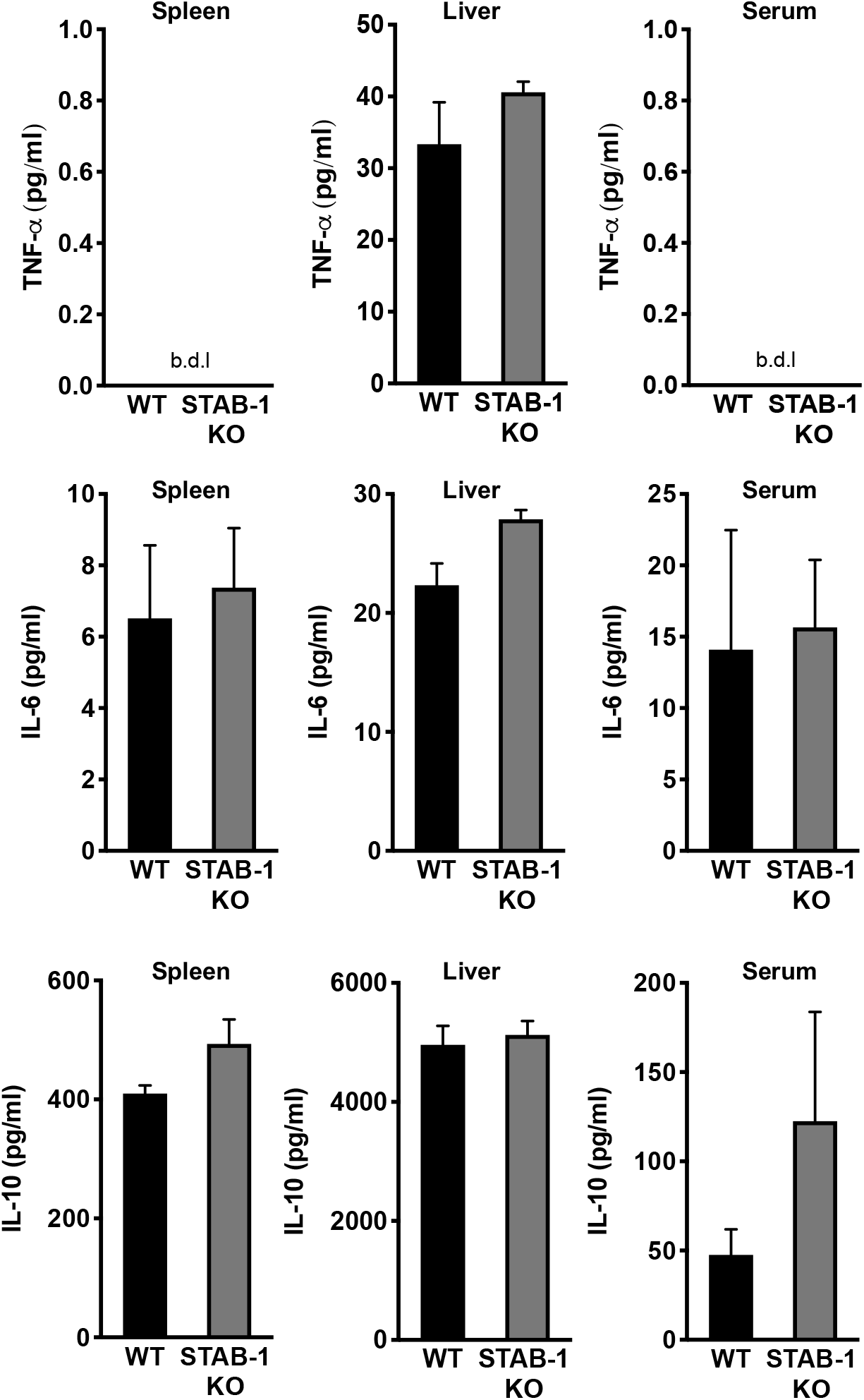
STAB-1 KO mice do not have significant defect on cytokine production. Cytokine production (TNF-α, IL-6, IL-10) in spleen, liver and serum from non-infected WT and STAB-1 KO mice was quantified by ELISA (n=3).

**Figure S2.**
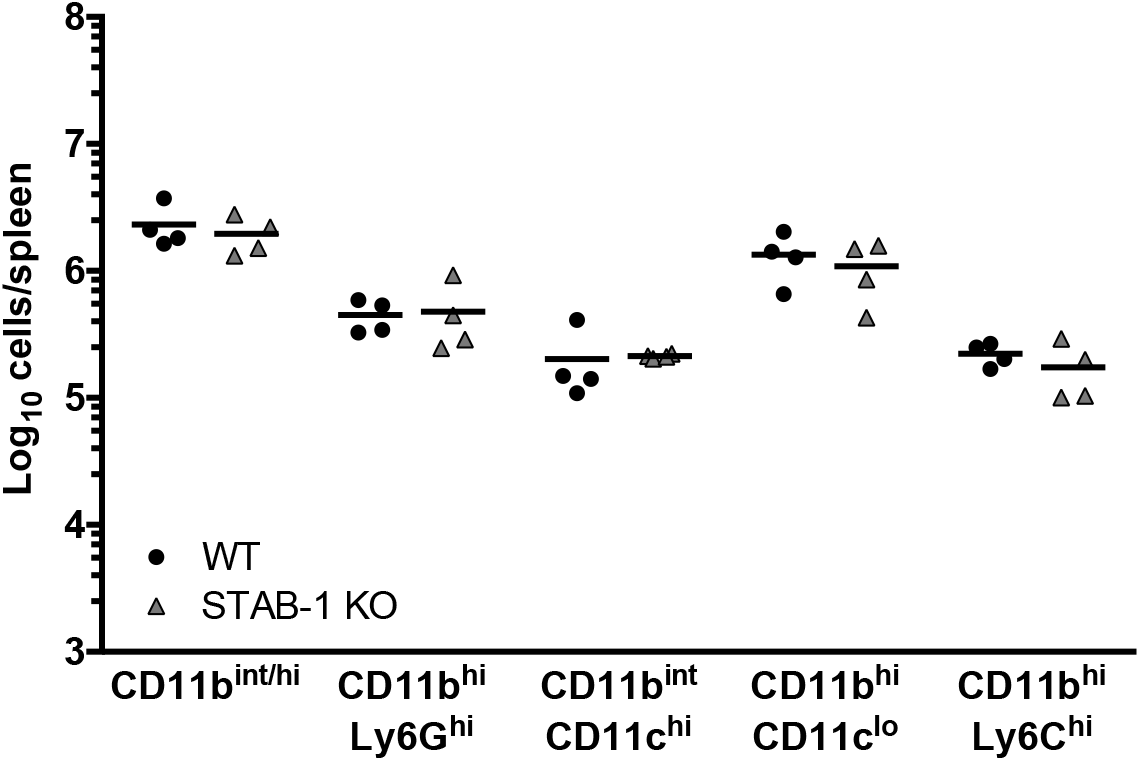
STAB-1 KO mice do not have significant defect on myeloid cell population. Spleen cells from non-infected WT and STAB-1 KO mice were isolated and analysed by flow cytometry to evaluate cell populations. Total numbers of myeloid cells (CD11b^int/hi^), neutrophils (CD11b^hi^Ly6G^hi^), dendritic cells (CD11b^int^CD11c^hi^), macrophages (CD11b^hi^CD11c^lo^) and inflammatory macrophages (CD11b^hi^Ly6C^hi^) are shown. Data are presented as scatter plots, with each animal represented by a dot and the mean indicated by a horizontal line.

**Table S1.**
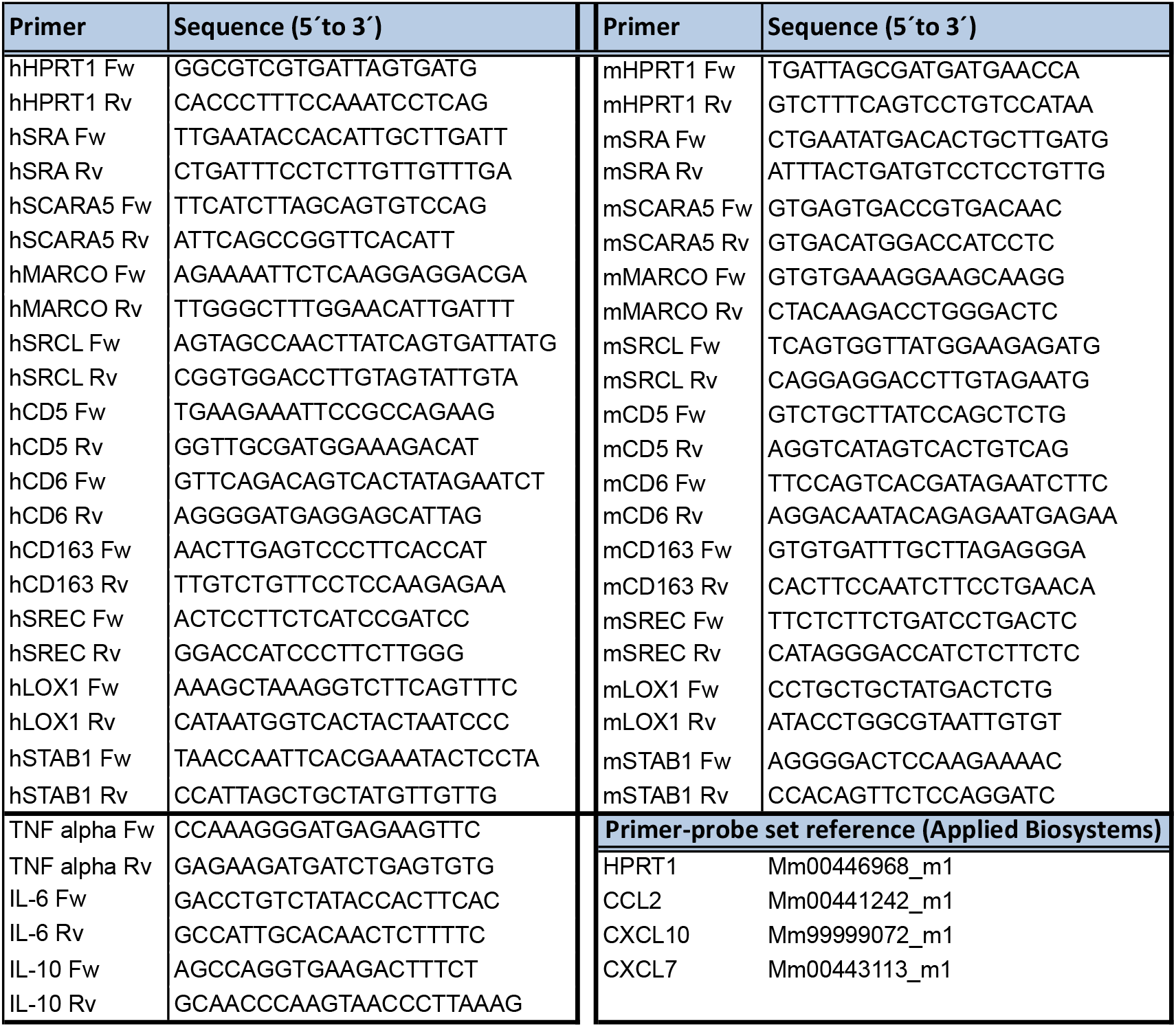
Priers

